# Visible-Light Optical Coherence Tomography Fibergraphy of the Tree Shrew Retinal Ganglion Cell Axon Bundles

**DOI:** 10.1101/2023.05.16.541062

**Authors:** David A. Miller, Marta Grannonico, Mingna Liu, Elise Savier, Kara McHaney, Alev Erisir, Peter A. Netland, Jianhua Cang, Xiaorong Liu, Hao F. Zhang

## Abstract

We seek to develop techniques for high-resolution imaging of the tree shrew retina for visualizing and parameterizing retinal ganglion cell (RGC) axon bundles in vivo. We applied visible-light optical coherence tomography fibergraphy (vis-OCTF) and temporal speckle averaging (TSA) to visualize individual RGC axon bundles in the tree shrew retina. For the first time, we quantified individual RGC bundle width, height, and cross-sectional area and applied vis-OCT angiography (vis-OCTA) to visualize the retinal microvasculature in tree shrews. Throughout the retina, as the distance from the optic nerve head (ONH) increased from 0.5 mm to 2.5 mm, bundle width increased by 30%, height decreased by 67%, and cross-sectional area decreased by 36%. We also showed that axon bundles become vertically elongated as they converge toward the ONH. Ex vivo confocal microscopy of retinal flat-mounts immunostained with Tuj1 confirmed our in vivo vis-OCTF findings.

## I. Introduction

Different animal models have been used to study the development and function of the visual system and eye diseases [1]. Mouse models are most commonly used given their genetic malleability [2], relatively short reproductive cycles, and low costs [1, 3]. However, mouse eyes lack key anatomical and functional similarities to human eyes, including a “fovea-like” region of high visual acuity (area centralis) [4], high cone density and color vision [3], and stratified retinal nerve fiber layer (RNFL) [5]. On the other hand, non-human primates [6] and larger animal models, such as dogs [7] and pigs [8], that share more of these features, are costly to house, have longer reproductive cycles, and face greater institutional and ethical restrictions [3], making them less ideal for large scale studies.

Tree shrews have emerged as a new eye model because they share greater similarities with the human retina than rodent models. For example, tree shrew eyes have been used to model refractive development [9], myopia [10], diabetic retinopathy [11], glaucoma [12], and central vision processing [13]. Noninvasive *in vivo* imaging techniques, such as optical coherence tomography (OCT) or scanning laser ophthalmoscopy (SLO), have been applied to examine the anatomical similarities and differences between tree shrew and human retinas [3, 12, 14]. Samuels *et al*. tracked the bulk RNFL thickness changes measured using near infrared (NIR) OCT in response to prolonged ocular hypertension (OHT) [12]. However, the NIR-OCT used in this study and others [3, 14] lacked the necessary axial resolution to clearly visualize the individual retinal ganglion cell (RGC) axon bundles that make up the RNFL. The size parameters of individual bundles serve as a more sensitive biomarker for RGC damage with glaucoma progression than bulk RNFL thickness change [15-17]. Therefore, we optimized a novel imaging technique, visible-light OCT fibergraphy (vis-OCTF) [15, 16], to non-invasively visualize individual RGC axon bundles and their surrounding microvasculature in the living tree shrew retina.

In OCT imaging, self-interference of the low-coherence source within the sample tissue generates a noise pattern known as speckle noise [18]. Speckles manifest as clusters of bright or dark pixels, which make resolving the boundaries between scattering structures difficult. This issue can be addressed by averaging multiple image acquisitions after decorrelating the speckle patterns through phase front modulation [19, 20], spatial averaging [21], or motions within the sample itself [22, 23]. Recently, Zhang *et al*. introduced temporal speckle averaging (TSA), which consists of capturing a series of OCT volumes over the same region of interest (ROI) in the retina [22]. Bulk tissue motions from respiratory and cardiac cycles and sub-cellular motions between volume acquisitions effectively decorrelate the speckle pattern. Thus, significant speckle reduction throughout the entire image volume can be achieved after careful registration and averaging of repeated image volumes [22, 23].

In this study, we tested whether combining vis-OCTF with TSA in the tree shrew retina could enhance the visualization of individual RGC axon bundles. First, we demonstrate methods for acquiring high-resolution vis-OCT images of the tree shrew retina using TSA [22, 23] and show the image quality improvements. Then, we demonstrate vis-OCT angiography (vis-OCTA), enhanced by TSA, to visualize the retinal microvasculature of the tree shrew retina. Next, we visualize individual bundle distribution *in vivo* using vis-OCTF and validate this distribution *ex vivo* using confocal microscopy. Lastly, we describe methods for quantifying single-bundle width, height, and cross-sectional area using vis-OCTF.

## II. Materials and Methods

### A. Animal Preparation

We imaged healthy 5–36-month-old northern tree shrews (*Tupaia belangeri*). Before imaging, all eyes were screened for cataracts by visually inspecting the cornea clarity. Animals with suitable eye quality were anesthetized using 5% isoflurane with supplemental oxygen at a flow rate of 1L/min. Tropicamide (1%; Henry Schein Animal Health) and phenylephrine (2.5%; Henry Schein Animal Health) eye drops were given to dilate the pupils and induce cycloplegia. During imaging, animals were maintained under anesthesia using 1-3% isoflurane with supplemental oxygen. Animals were kept warm using an infrared heat lamp, and polyvinyl alcohol artificial tears (1.4%; Rugby Laboratories, Inc., Hempstead, NY) were applied between acquisitions to prevent corneal dehydration. After imaging, animals were placed on a heating pad and monitored until alert and active.

### B. Visible-Light Optical Coherence Tomography

Tree shrews were imaged using a commercial small animal vis-OCT retinal imaging system (Halo 100; Opticent Health, Evanston, IL), as described previously [15, 16]. The system, illustrated in Fig. 1a, used a filtered supercontinuum light source to deliver visible light to a 2 × 2 fiber coupler. The reference arm consisted of a polarization controller, glass plates for dispersion compensation, and a mirror on a motorized translation stage. The sample arm consisted of a pair of galvanometer scan mirrors and a 3:1 Keplerian telescope. The incident power delivered to the cornea was 1 mW. Returned interference fringes were detected using a commercial spectrometer (Blizzard SR; Opticent Health) with a spectral detection range from 508 nm to 613 nm. Interference fringes were acquired with an A-line rate of 75 kHz (an integration time of 12.6 μs). The axial resolution of the system was 1.3 μm in the retina. The lateral resolution was 4.5 μm at the center of the field of view and 8.7 μm at 560 μm from center as approximated by the Norton tree shrew eye model [9]. The total image volume was approximately 1.12 mm (x) × 1.12 mm (y) × 1.5 mm (z).

**Fig. 1.**
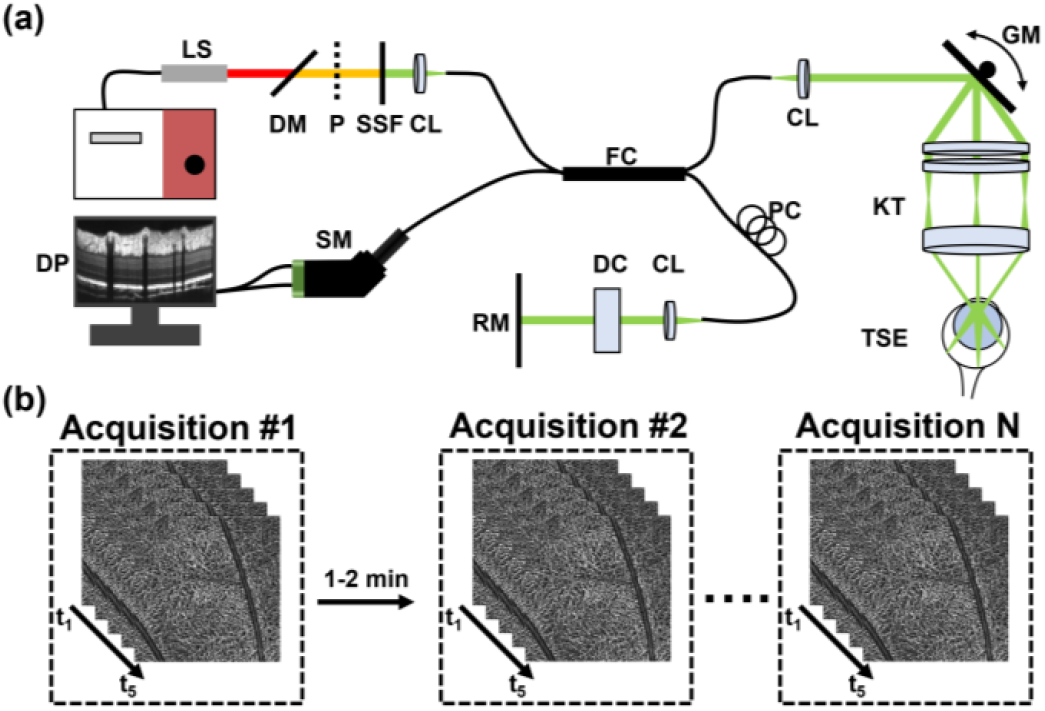
System schematic and image acquisition protocol. (a) Schematic of the small-animal vis-OCT system for tree shrew imaging. CL: collimating lens; DC: dispersion compensation; DP: data processing; DM: dichroic mirror; FC: 2 × 2 fiber coupler; GM: galvanometer scan mirrors; KT: Keplerian telescope; LS: supercontinuum light source; P: polarizer; PC: polarization controller; RM: reference mirror; SM: spectrometer; SSF: spectral shaping filter; TSE: tree shrew eye. (b) Image acquisition flow for short and long TSA datasets. Single TSA consists of five repeated vis-OCTA volumes. Long TSA consists of multiple acquisitions taken 1-2 min apart.

For each acquisition, we followed a TSA acquisition protocol to reduce speckle noise [22, 23]. As demonstrated by Fig. 1b, a single acquisition consisted of five repeated vis-OCTA volumes without delay between repeated volumes. A single vis-OCTA volume consisted of 512 A-lines/B-scan × 512 B-scans with each B-scan repeated twice to identify blood flow contrast. A single acquisition took ∼35 seconds to complete. The repeated volumes were registered and averaged to generate a TSA dataset, as detailed in the following section. For higher quality reconstructions, multiple TSA datasets were acquired at the same location, as depicted in Fig. 1b, and averaged following the same procedure to generate a high-quality TSA dataset.

### C. Temporal Speckle Averaging

Structural vis-OCT volumes were processed using background subtraction, automatic sub-band dispersion compensation (detailed in Section 2D), *k*-space resampling, and Fourier transformation. Angiograms were generated from repeated B-scans following our previous protocol [24].

Fig. 2 shows the TSA processing steps for registering and averaging repeated vis-OCT volumes to an optimal reference volume. This optimal reference volume maximizes the number of volumes that can be utilized with minimal alignment error. The optimal reference volume is selected, and each volume is aligned to it as follows: *<u>Step 1</u>*: Enface angiograms are extracted from each repeated vis-OCTA volume. The angiogram from the first vis-OCTA volume is selected as the first reference image. *<u>Step 2</u>*: Angiograms are coarsely aligned to the reference angiogram using rigid registration transformations (Step 2a) followed by fine alignment with non-rigid image transformation using a gradient descent method (Step 2b). The alignment quality after enface registration is assessed using a structural similarity index (SSI) and stored in an alignment quality matrix. Step 2 is repeated with each angiogram as the reference image. *<u>Step 3</u>*: The optimal reference volume is determined from the alignment quality matrix by selecting the reference volume with the highest mean SSI across all other images. In Fig. 2, the optimal volume is indicated by the dashed black box. Optimal rigid and non-rigid image transformations are applied to each depth position of each vis-OCT and vis-OCTA volume. *<u>Step 4</u>*: Time-series B-scans are extracted from the laterally registered volumes for depth registration. *<u>Step 5</u>*: Each B-scan is up-sampled six times in both dimensions and registered to the B-scan from the reference volume using rigid registration. *<u>Step 6</u>*: Each A-line from the registered B-scan is extracted (Step 6a) and aligned to the reference A-line by cross correlation (Step 6b). *<u>Step 7</u>*: Registered B-scans are averaged and down-sampled to generate averaged vis-OCT and vis-OCTA volumes.

**Fig. 2.**
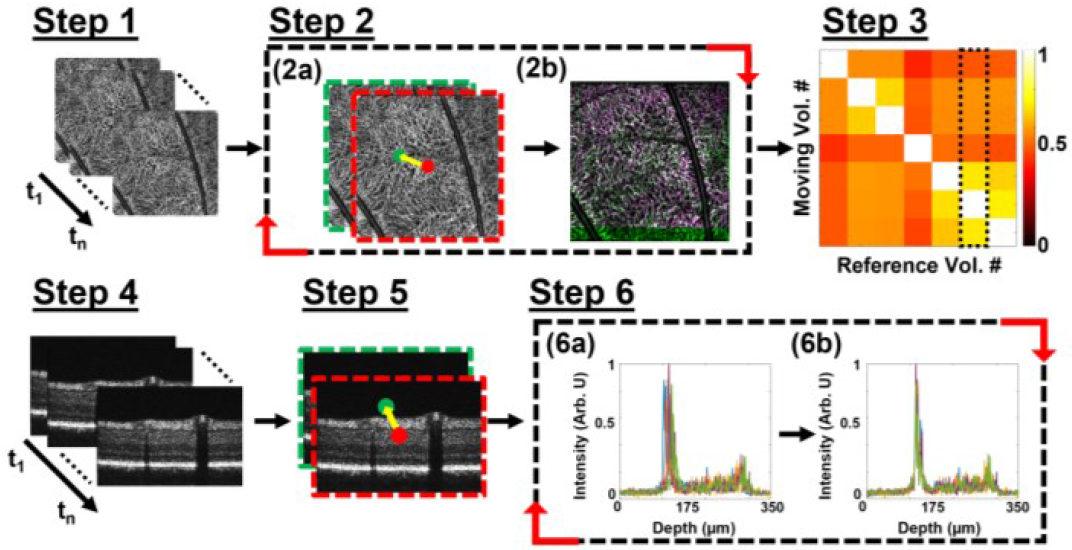
TSA image registration pipeline. Step 1: time-series angiogram extracted from repeated vis-OCTA volumes. Step 2: (2a) rigid registration followed by (2b) non-rigid registration of moving image to reference image. Step 3: Optimal reference volume selected from alignment quality matrix (dashed black box). Step 4: time-series B-scan extracted from each volume. Step 5: Rigid registration of each B-scan to reference B-scan. Step 6: (6a) Extracted A-lines and (6b) A-lines after registration by cross-correlation.

High-quality TSA datasets, with >5 averages, were generated by capturing multiple five-volume TSA datasets from the same location with 1–2-minute intervals between acquisitions, as depicted in Fig. 1b. These acquisitions were then registered and averaged following the same TSA pipeline in Fig. 2.

### D. Automatic Sub-band Dispersion Compensation

Dispersion mismatch between the sample and reference arms of an OCT system leads to wavenumber (*k*) dependent non-linear phase changes in the OCT signal. Such non-linear phase changes distort the point spread function (PSF) in the spatial domain, thus reducing the axial resolution of the image [25]. This effect can be minimized by adding dispersion compensation glass to the reference arm to match the optical pathlength until the PSF is fully optimized. To account for remaining dispersion mismatch introduced by the sample, digital dispersion compensation can be performed to numerically cancel the *k* dependent non-linear phase. The complex OCT signal can be expressed as

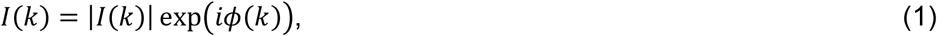

where |*I*(*k*)| is the amplitude and *ϕ*(*k*) is the phase. To correct for dispersion, the phase of *I*(*k*) is modified by including a phase correction term, *ϕ*_*c*_(*k*),

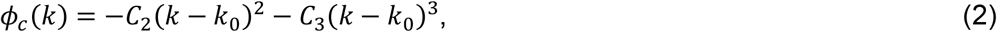

where *k*_0_ represents the center wavenumber, and *C*_2_ and *C*_3_ represent the phase correction coefficients [25]. The dispersion-corrected complex OCT signal can be expressed as

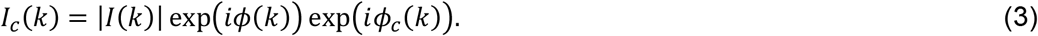

We implemented an automated approach using spectroscopic analysis to select the optimal phase correction coefficients for any OCT image. First, the phase correction coefficients are initialized to zero, and short-time Fourier transform (STFT) is applied to *I*_*c*_(*k*) to generate spectroscopic B-scans. The spectroscopic B-scans are then up-sampled in the axial dimension, and the absolute spectral shift between spectral B-scans is calculated. The absolute spectral shift is defined as

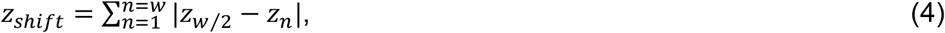

where *w* is the total number of STFT windows. An optimization routine is then used to update the phase correction coefficients until *Z*_*shift*_ is minimized.

Unlike full-band dispersion compensation methods that optimize metrics such as sharpness [25] or entropy [26] within the reconstructed image, our sub-band dispersion compensation method directly optimizes the spectral separations that give rise to dispersion. Additionally, this technique requires no calibration, making it suitable for highly dispersive samples.

### E. Fibergram Processing and Analysis

Fibergram images were generated as described previously [15, 16]. Briefly, vis-OCT images were acquired with retina positioned at a uniform depth throughout the volume to maximize reflectance from the RGC axon bundles. After TSA processing, fibergrams were extracted by segmenting the RNFL and taking the mean intensity projection along the axial dimension. The width of individual RGC axon bundles was measured from fibergram images by manually marking the center axis of the bundle. We then reconstructed the intensity profile orthogonal to the center axis and normalized the curve between 0 and 1. The bundle width was recorded as the intensity profile width at 1/e^2^. The height of individual bundles was measured from B-scans that were resampled orthogonal to the previously marked center axis of the bundles. Bundle height was similarly measured by extracting the axial intensity profile of the bundle and recording the value at 1/e^2^. The cross-sectional area of the bundles (*A*_*b*_) was approximated by the area of an ellipse as

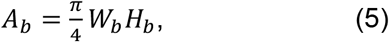

where *W*_*b*_ is bundle width and *H*_*b*_ is bundle height [17].

### F. Image Quality Metrics

To demonstrate the image quality improvement after performing TSA, we quantified the background noise level (*σ*_*bg*_), signal-to-noise ratio (*SNR*), and contrast-to-noise ratio (*CNR*). The background noise level was defined as the standard deviation of the image intensity in the vitreous just above the RNFL. SNR describes the relationship between the background pixel uncertainty and the mean signal value, defined as

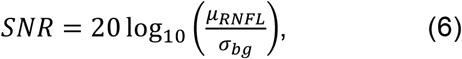

where *μ*_*RNFL*_ is the mean image intensity within the RNFL.

*CNR* describes the image contrast with respect to pixel uncertainty. Thus, in images with higher *CNR*, portions of the image containing signal from the retina are more distinguishable from the surrounding background. We defined *CNR* as

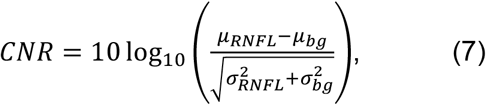

where *μ*_*bg*_ is the mean background image intensity and *σ*_*RNFL*_ is the standard deviation of the RNFL image intensity.

### G. Immunohistochemistry and Confocal Microscopy

After acquiring vis-OCT data, tree shrews were euthanized with euthasol (3.9mg/mL pentobarbital, 0.5mg/mL phenytoin sodium; Virbac ANADA, # 200-071) and perfused with paraformaldehyde (PFA) (4%; ChemCruz, sc-281692). Retinas were dissected, post-fixed in PFA for 30 minutes, washed with phosphate buffered saline (PBS) containing Triton-X detergent (PBST, 0.5% Triton X-100), and blocked for 2 hours at room temperature in blocking buffer (2.5% BSA, 5% normal donkey serum, and 0.5% Triton X-100 (Sigma-Aldrich)). Samples were incubated with the primary antibody, mouse anti-Tuj-1 (1:200; a gift from Anthony J. Spano), overnight at 4°C. The secondary antibody, donkey anti-mouse immunoglobulin G, conjugated to Alexa Fluor 488 dye (Invitrogen A-21206, RRID: AB_2535792), was diluted at 1:1000 in blocking buffer and incubated overnight at 4°C. The whole-mount retinas were cut into four quadrants: temporal (T), nasal (N), inferior (I), and superior (S), flat-mounted, and cover-slipped with Vectashield mounting medium (Vector Laboratories Inc) [15, 27]. Confocal microscopy was performed using the 3D Z-stack mode on a Zeiss LSM 800 microscope (Carl Zeiss AG) [15, 28]. To capture the central region of the flat-mounted three shrew retina, tiles were imaged to cover a total volume of 7.00mm (x) × 7.0000 mm (y) × 90 μm (z) at a pixel size of 1.24 μm/pixel. Z-stack slices were then projected to create 2-D enface confocal microscopy images.

### H. Statistical Analysis

The one-way analysis of variance (ANOVA) test was used to compare the image quality metrics and bundle size parameters for each case. Comparison between groups was performed using a post-hoc Tukey test. A significance level of 0.05 was used for all statistical comparisons. All results were reported as mean ± standard deviation.

## III. Results

### A. TSA-enhanced Tree Shrew Retinal Imaging

We assessed TSA’s ability to resolve the finer details in the tree shrew retina that are normally obscured by speckle noise. Figs. 3a-3b show typical mean-intensity-projection enface (Fig. 3a) and B-scan images reconstructed along the red dashed lines (Fig. 3b) of the tree shrew retina without averaging (left), after 10 volume averages (middle), and after 70 volume averages (right). We observed a reduction in speckle noise and increased image detail as the volume averages increased. For example, in Fig. 3a, finer details, such as the small microcapillaries indicated by the green arrows, became visible with increased volume averages. Similarly, in Fig. 3b, increased volume averages revealed the boundaries of individual RGC axon bundles, a dark band in the inner plexiform layer (IPL), and a sharper outer retina structure. Video 1 (see Supplementary Materials) shows a B-scan fly-through of the 70 times averaged image, further highlighting improved quality throughout the volume. Video 2 (see Supplementary Materials) shows a fly-through of enface images extracted from each depth location.

**Fig. 3.**
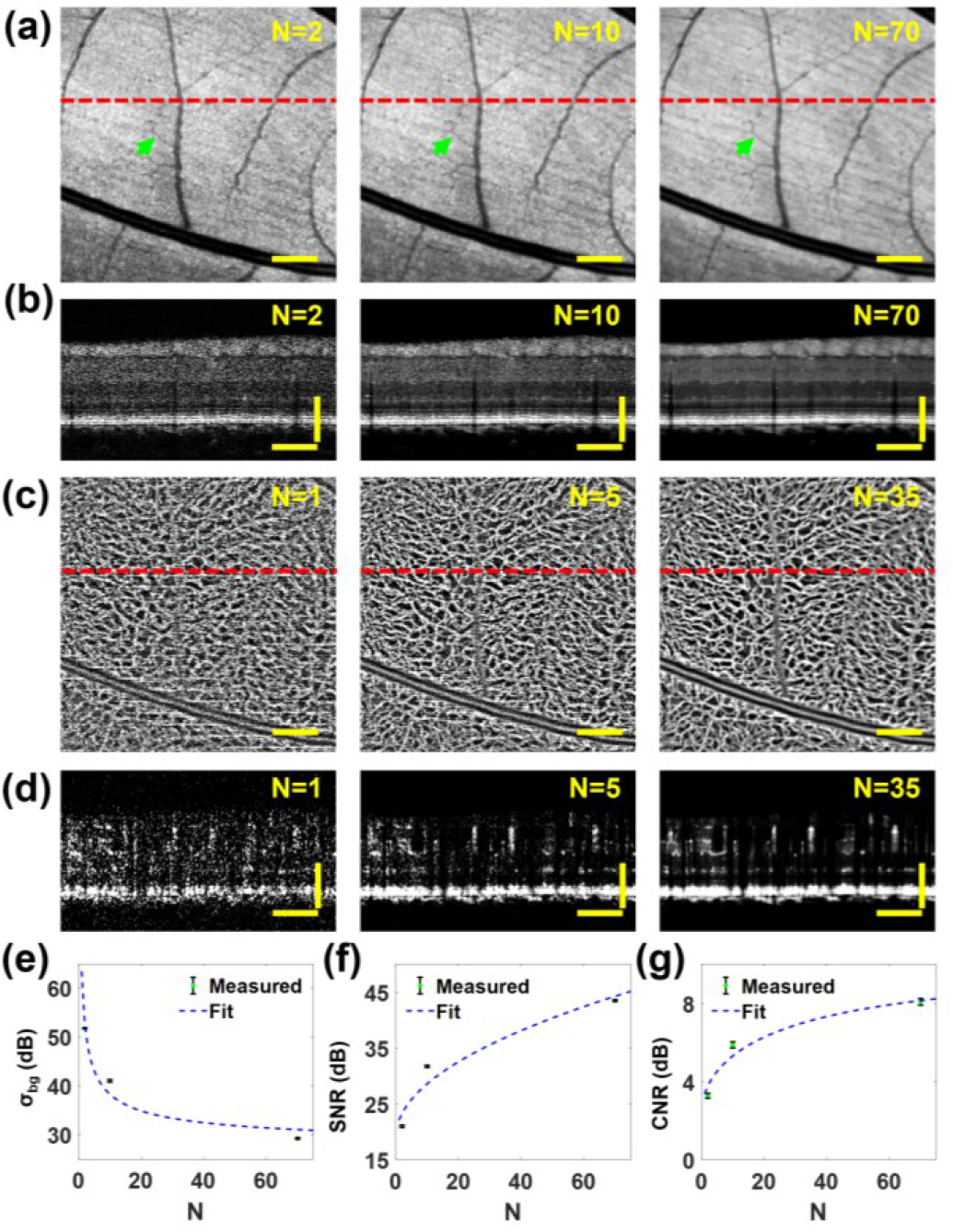
TSA image quality comparison with increased number of averages. (a) En-face image without volume averaging (left), 10 averages (middle), and 70 volume averages (right). (b) B-scan images without volume averaging (left), 10 volume averages (middle), and 70 volume averages (right). (c) Maximum-intensity-projection angiogram without volume averaging (left), five volume averages (middle), and 35 volume averages (right). (d) Vis-OCTA B-scans without volume averaging (left), with five volume averages (middle), and 35 volume averages (right). (e) Changing σ_bg_ with *N*. (f) Changing SNR with *N*. (g) Changing CNR with *N*. N: number of volume averages; M: number of decorrelated speckle patterns; Scale bars: 150 μm.

We applied vis-OCTA to determine if the increased volumetric averaging also improved vis-OCTA image quality. Fig. 3c shows a reconstructed angiogram without volumetric averaging (left), after 5 volume averages (middle), and after 35 volume averages (right). Compared to the unaveraged angiogram in Fig. 3c, the angiogram after TSA presents higher contrast between blood vessels and the image background, resulting in a more defined blood vessel structure. Similarly, the vis-OCTA B-scans with more averages in Fig. 3d, reconstructed along the dashed red lines in Fig. 3c, show reduced noise resulting in sharper depth-resolved blood vessels.

We quantified *σ*_*bg*_, *SNR*, and *CNR* in 150 vis-OCT B-scan images with 2, 10, and 70 averages and plotted them in Figs. 3e-3g. As the number of volumetric averages increased, *σ*_*bg*_ decreased from 51.8±0.2 dB with 2 averages to 41.1±0.2 dB with 10 averages, and 29.3±0.2 dB with 70 averages, as shown in Fig. 3e. This decrease in *σ*_*bg*_ is proportional to 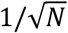 (dashed line in Fig. 3e), the theoretical decrease in *σ*_*bg*_ with *N. SNR* increased from 20.9±0.2 dB with 2 averages to 31.7±0.2 dB with 10 averages, and 43.5±0.2 dB with 70 averages. This trend, depicted in Fig. 3f, is consistent with the theoretical *SNR* improvement with increased volumetric averages, which follows 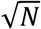, as indicated by the dashed line in Fig. 3f. Lastly, the *CNR* plot in Fig. 3g shows an increase from 3.3±0.1 dB with 2 averages to 5.8±0.2 dB with 10 averages to 8.1±0.2 dB with 70 averages, following the relationship 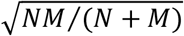, where *M* is the number of uncorrelated speckle patterns, indicated by dashed line in Fig. 3g.

Fig. 4 shows high-quality TSA imaging with 30 volume averages near the optic nerve head (ONH). A fly-through of all the B-scans in this volume is shown in Video 3 (see Supplementary Materials), and a fly-through of enface images extracted from each depth location is shown in Video 4 (see Supplementary Materials). Fig. 4a shows a resampled circumpapillary B-scan [16] reconstructed along the 350 μm radius red circle shown in the fundus image on the left of Fig. 4a. The blue arrows in Fig. 4a indicate the leftmost A-line in the B-scan and the green arrow indicates the direction along which the B-scan was reconstructed. The B-scan in Fig. 4b is a magnified view of the dashed yellow box in Fig. 4a with the inner retinal layers labeled. In Fig. 4a, we again observed individual axon bundles in the RNFL and a dark band within the IPL. Fig. 4c is a magnified view of the dashed box in Fig. 4b with the outer retinal layers labeled. In the outer retina, we observed a thin outer nuclear layer (ONL), which included a bright sub-band near the top of the layer.

**Fig. 4.**
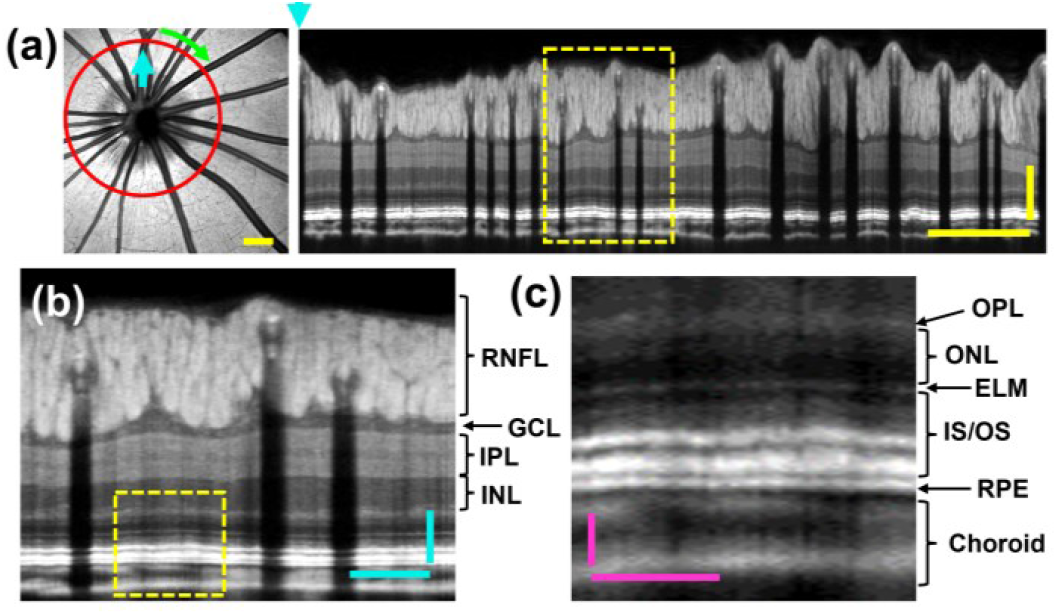
ONH TSA image with 30 volume averages. (a) En-face (left) and resampled circumpapillary B-scan reconstructed along the red circle. Scale bars: 150 μm. (b) Magnified view of dashed box in (a) with inner retinal layers labeled. Scale bars: 50 μm. GCL: ganglion cell layer; INL: inner nuclear layer (c) Magnified view of dashed yellow box in (b) with outer retinal layers labeled. Scale bars: 25 μm. OPL: outer plexiform layer; ELM: external limiting membrane; IS/OS: inner segment/outer segment junction; RPE: retinal pigment epithelium.

We generated large FOV images by montaging multiple acquisitions together after TSA processing. Fig. 5a shows a montaged fibergram. This montage shows how the RGC axon bundles are distributed throughout the retina in the tree shrew. Fig. 5b shows a montaged angiogram of the tree shrew, revealing high microvasculature density throughout the retina.

**Fig. 5.**
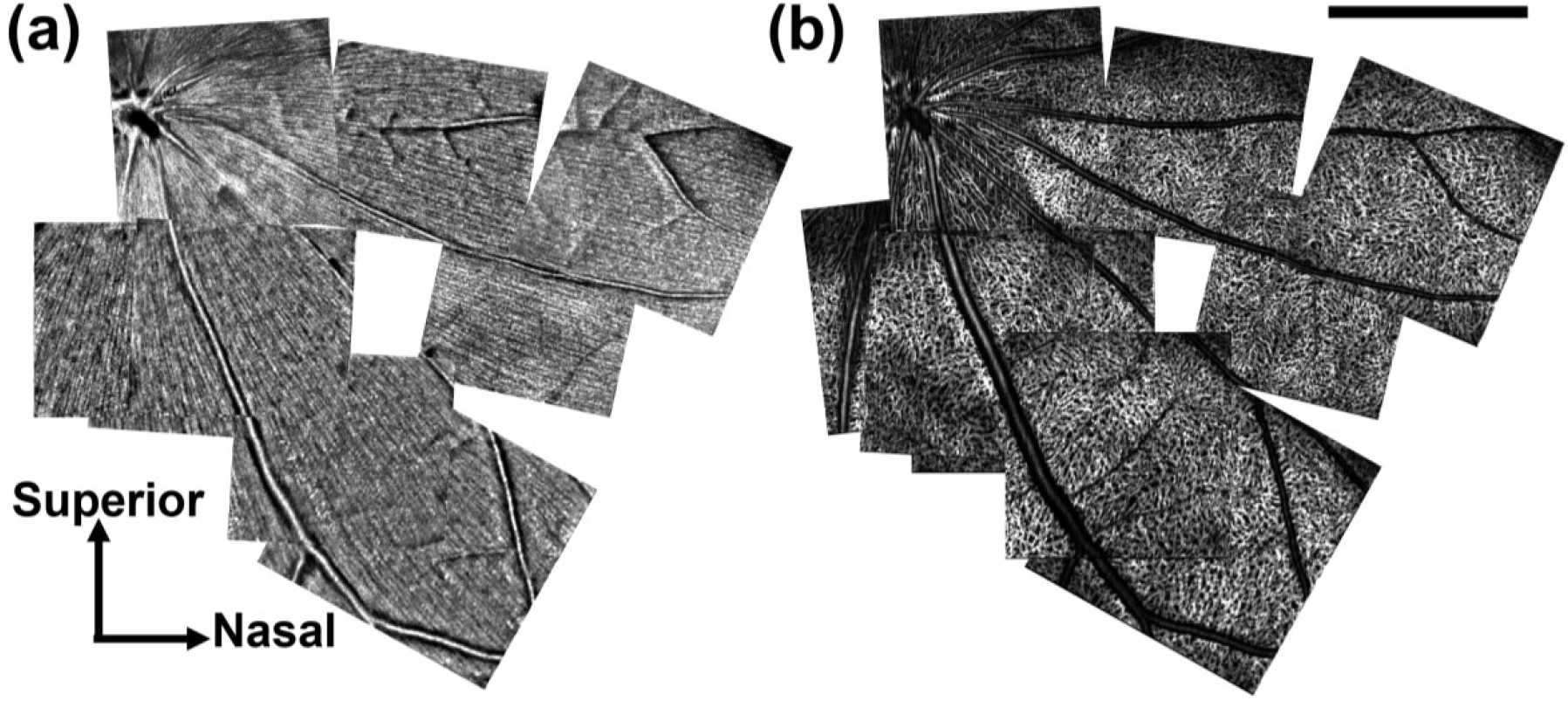
Montage images of Tree Shrew Retina. (a) Montaged fibergram. (b) Montaged angiogram. Scale bar: 1 mm.

### B. RGC Axon Bundle Organization

We combined TSA with vis-OCTF to further characterize the RGC axon bundle structure at a resolution that was not previously achieved in the living tree shrew retina. Fig. 6 compares *in-vivo* TSA fibergram images and *ex-vivo* confocal microscopy images. The montaged vis-OCT fibergram (Fig. 6a) shows the large FOV map of the RGC axon bundle organization *in vivo*. We observed a high density of RGC axon bundles throughout most of the tree shrew retina. Fig. 6b shows a magnified fibergram within the region highlighted by the red dashed box in Fig. 6a and Fig. 6c shows the corresponding confocal microscopy image from the central retina. Figs. 6d and 6e respectively show the magnified views of vis-OCT fibergram and confocal microscopy images from the regions highlighted by the blue dashed box in Figs. 6a, which is ∼2 mm from the ONH. Side-by-side comparison of the *in vivo* and *ex vivo* images revealed identical bundle structures. The width of the axon bundles appears to increase moving outward from the ONH.

**Fig. 6.**
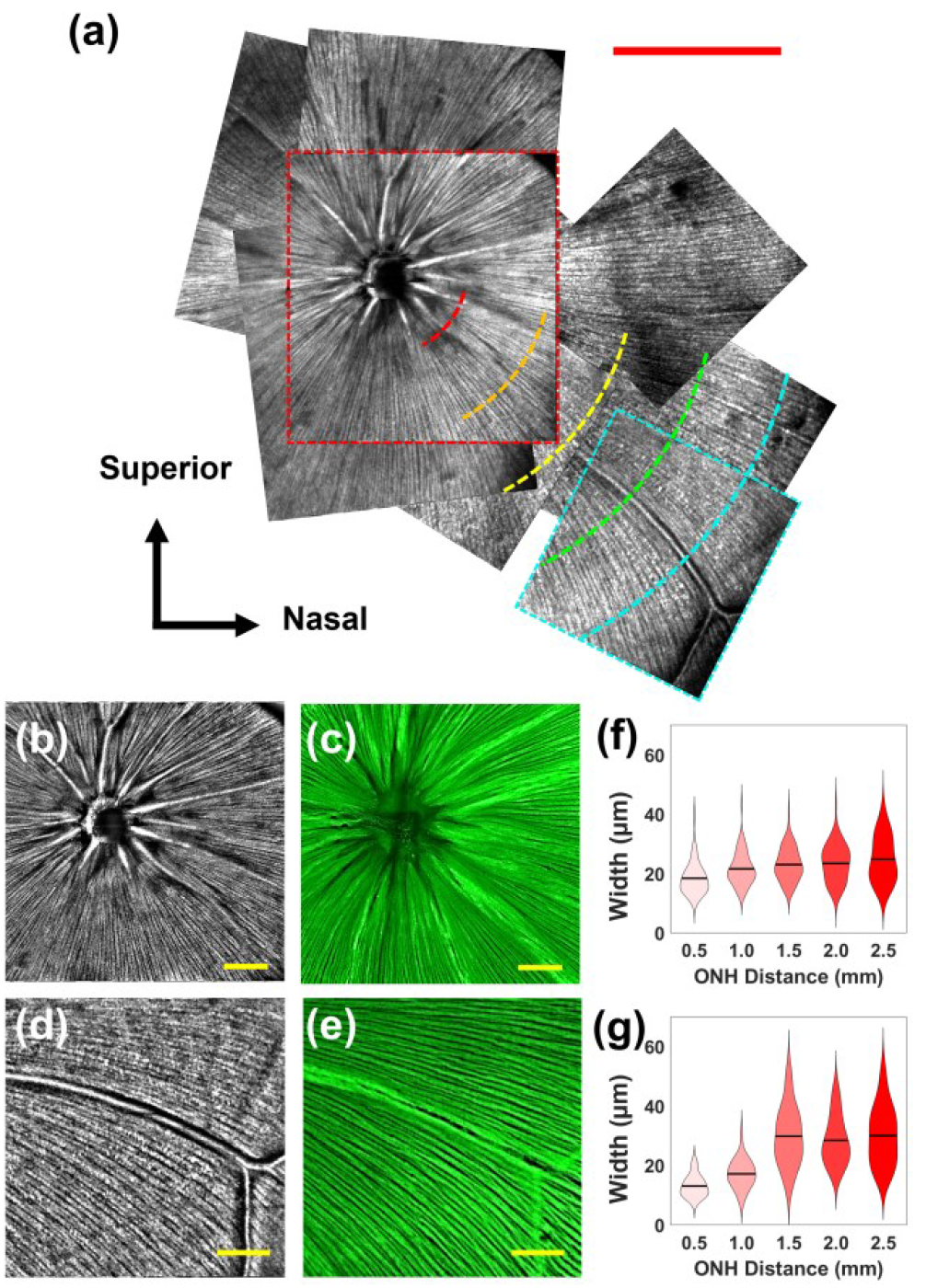
Vis-OCT fibergram and RGC bundle width analysis. (a) Montaged vis-OCT fibergram. (b) Magnified view of the red dashed box in (a). (c) Confocal microscopy image of retina immunolabeled with Tuj1 for RGC axon bundles acquired from the same location as (b). (d) Magnified view of blue dashed box in (a). (e) Confocal microscopy image of retina immunolabeled with Tuj1 for RGC axon bundles acquired from the same location as (d). (f) Varying RGC bundle width with respect to distance from the ONH. (g) Varying RGC bundle width with respect to distance from ONH within the 45° arc patterns as highlighted by the dashed lines in (a). Scale bars: red = 1 mm, yellow = 250 μm.

We measured the width of individual RGC axon bundles throughout the montaged tree shrew retina. Measurements were recorded at 0.5 mm intervals up to 2.5 mm from the ONH along concentric circular paths, as shown in Fig. 6a. Recorded RGC bundle widths with respect to the distance from the ONH are plotted in Fig. 6f. Bundle width was 19.8±4.3 μm (n=48) at 0.5 mm from the ONH, 21.5±5.4 μm (n=131) at 1 mm, 23.1±5.5 μm (n=95) at 1.5 mm, 23.5±6.3 μm (n=75) at 2 mm and 24.9±7.5 μm (n=51) at 2.5 mm. A post-hoc Tukey test showed a significant difference between bundle width at 0.5 mm and all groups, and a significant difference in bundle width between 1.0 mm and 2.5 mm. The standard deviation of the bundle width also increased with distance from the ONH.

Next, we compared the width of continuous axon bundles within a 45° sector, as indicated by the yellow dashed lines in Fig. 6a. Within this 45° sector, we observed an increase in axon bundle width that plateaued near 1.5 mm from the ONH, as depicted in Fig. 6g. Bundle width was 13.1±3.6 μm (n=17) at 0.5 mm from the ONH, 17.2±4.7 μm (n=30) at 1 mm, 29.9±9.8 μm (n=27) at 1.5 mm, 28.5±8.1 μm (n=44) at 2 mm and 30.1±9.7 μm (n=51) at 2.5 mm. A post-hoc Tukey test showed a significant difference between bundle width at 0.5 mm and 1.5 mm, 0.5 mm and 2.0 mm, 0.5 mm and 2.5 mm, 1.0 mm and 1.5 mm, 1.0 mm and 2.0 mm, and 1.0 mm and 2.5 mm.

### C. RGC Axon Bundle Cross-Sectional Structure

Taking advantage of the high-resolution axon bundle visualization, we extracted the resampled vis-OCT B-scans from the montaged fibergram (Fig. 6a) to investigate the changing RGC bundle cross-sectional morphology from the ONH to the periphery. Figs. 7a-7e depict the B-scans reconstructed along the dashed arc paths at 0.5 mm (Fig. 7a), 1.0 mm (Fig. 7b), 1.5 mm (Fig. 7c), 2.0 mm (Fig. 7d), and 2.5 mm (Fig. 7e) from the ONH. Figs. 7f-7j show the magnified view of the dashed boxes in Figs. 7a-e. As the distance from the ONH increased, we observed an overall reduction in axon bundle height and a change in the organization of individual bundles. For example, the magnified view at 0.5 mm from the ONH in Fig. 7f shows stratified and vertically elongated bundles, as indicated by green arrows. Moving outward from the ONH to the periphery, the bundles spread out, forming a monolayer of round axon bundle cross sections, as shown in Fig. 7j.

**Fig. 7.**
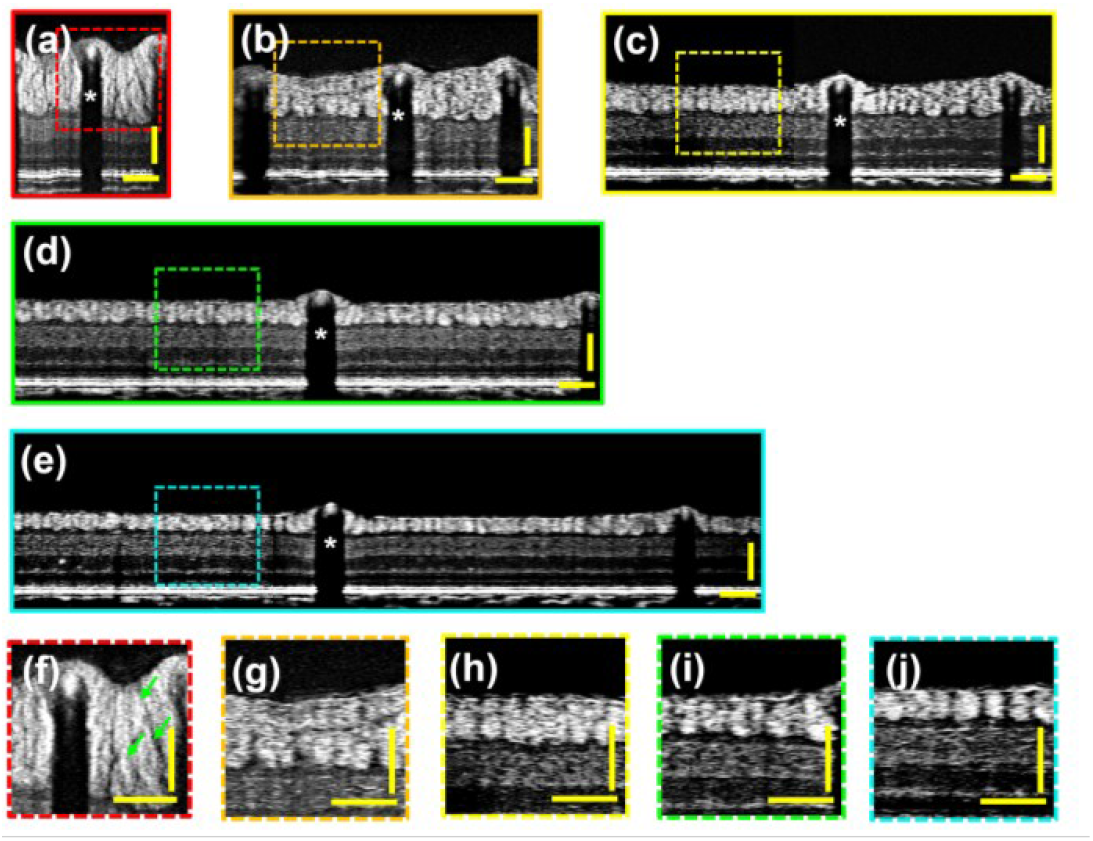
Resampled B-scans from tree shrew retina at different distances from the ONH highlighting RGC bundle cross-sectional structure. Resampled circumpapillary B-scans reconstructed along arc traces in Fig. 6a at (a) 0.5 mm, (b) 1.0 mm, (c) 1.5 mm, (d) 2.0 mm, and (e) 2.5 mm from ONH. White stars indicate the same blood vessel in (a-e). (f)-(j) are magnified views of the regions highlighted by the dashed boxes in (a)-(e). The green arrows in (f) indicate stacked RGC bundles. Scale bars: black = 1 mm, yellow = 100 μm.

We measured the heights of individual RGC bundles in the fibergram images at 0.5 mm distance increments from the ONH. The bundle heights in Fig. 8a are 89.6±31.1 μm (n=48) at 0.5 mm, 63.0±24.0 μm (n=131) at 1.0 mm, 56.7±16.7 μm (n=95) at 1.5 mm, 44.4±10.7 μm (n=75) at 2.0 mm, and 35.2±6.3 μm (n=51) at 2.5 mm. A post-hoc Tukey test showed a significant difference between bundle height at 0.5 mm and all groups, 1.0 mm and 2.0 mm, and 1.0 mm and 2.5 mm. In addition to the overall bundle height decrease, we observed a decrease in the standard deviation.

**Fig. 8.**
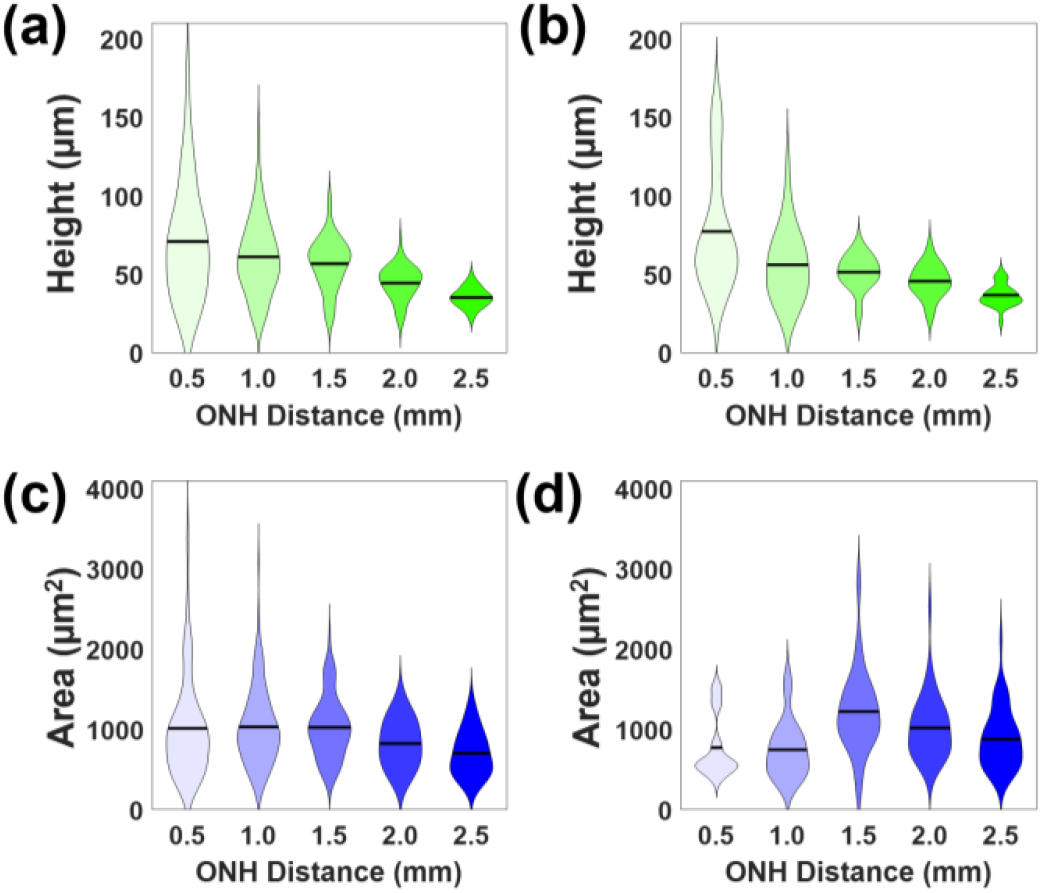
RGC bundle cross-sectional size analysis. (a) Bundle height changes with respect to ONH distance in the whole retina. (b) Bundle height measurements acquired along arc paths in Fig. 6a as a function of distance from the ONH. (c) Cross-sectional bundle area measurements from the whole retina. (d) Cross-sectional bundle area measurements acquired along arc paths.

Next, we tracked the change in height within the same continuous 45° sector as Fig. 6a. Similar to the bundle height change in the whole retina, we observed a reduction in height for individual bundles when the distance increased from the ONH. Similarly, standard deviation was the highest near the ONH, where bundles formed vertically elongated cross-sections. Bundle height was 77.3±37.8 μm (n=17) at 0.5 mm, 56.0±22.8 μm (n=30) at 1 mm, 51.4±10.7 μm (n=27) at 1.5 mm, 45.6±10.6 μm (n=44) at 2 mm and 36.7±7.0 μm (n=51) at 2.5 mm. A post-hoc Tukey test showed a significant difference between bundle height at 0.5 mm and all groups, 1.0 mm and 2.5 mm, and 1.5 mm and 2.5 mm. These observations are plotted in Fig. 8b.

Lastly, we calculated the bundle cross-sectional area with respect to the distance from the ONH. We observed an overall decrease in bundle cross-sectional area with increasing distance from the ONH, as shown in Fig. 8c. The bundle area was 1400±602 μm^2^ (n=48) at 0.5 mm, 1100±468 μm^2^ (n=131) at 1.0 mm, 1000±389 μm^2^ (n=95) at 1.5 mm, 820±300 μm^2^ (n=75) at 2.0 mm, and 700±280 μm^2^ (n=51) at 2.5 mm. A post-hoc Tukey test showed a significant difference between bundle area at 0.5 mm and 2.5 mm, 1.0 mm and 2.0 mm, 1.0 mm and 2.5 mm, 1.5 mm and 2.0 mm, and 1.5 mm and 2.5 mm. The cross-sectional area standard deviation also decreases as the distance from the ONH increases.

Within the continuous 45° sector, we observed greater variation in bundle cross-sectional area. As shown in Fig. 8d, the bundle area was 771±390 μm^2^ (n=17) at 0.5 mm, 745±353 μm^2^ (n=30) at 1.0 mm, 1221±510 μm^2^ (n=27) at 1.5 mm, 1017±389 μm^2^ (n=44) at 2.0 mm, and 876±368 μm^2^ (n=51) at 2.5 mm. A post-hoc Tukey test showed a significant difference between bundle area at 0.5 mm and 1.5 mm, 1.0 mm and 1.5 mm, 1.0 mm and 2.0 mm, and 1.5 mm and 2.5 mm.

## IV. Discussion

In this study, we presented, for the first time, the visualization and analysis of individual RGC axon bundles in tree shrew retinas *in vivo*. We successfully imaged the boundaries of individual RGC axon bundles in fibergram and B-scan images by combining vis-OCTF with TSA. Such visualization enabled the quantification of single-bundle width, height, and cross-sectional area. Additionally, vis-OCTA combined with TSA revealed the microvasculature structure of the tree shrew retina, which has not been reported previously. Further, we validated the *in vivo* bundle distribution using *ex vivo* confocal microscopy images of the same retina immunolabeled by Tuj1 for RGC axons. Applying these acquisition, processing, and analysis techniques provides a foundation for future studies using the tree shrew animal model to study optic neuropathies with greater similarities to human pathology.

An optimized TSA processing algorithm was applied to reduce speckle noise and reveal the fine details of the tree shrew retina. Our results showed significantly reduced *σ*_*bg*_ and increased *SNR* and *CNR* by averaging a larger number of vis-OCT volumes. The results indicated 10 volume averages, acquired over ∼35 seconds, was adequate for distinguishing the boundaries of individual RGC axon bundles in the tree shrew retina. As the number of volume averages increased beyond 10 averages, more anatomical features became visible, including a dark band separating the top and bottom portions of the IPL. However, these acquisitions required up to 25 minutes to collect enough volumes, which is impractical for studies requiring a larger view of the retina, such as ours. Thus, we found 10 volume averages optimal for imaging a large retinal area while providing enough details to identify individual bundles.

Previous TSA studies demonstrated cellular-level resolution imaging in rodent eyes [22, 23]. For example, Zhang *et al*. demonstrated that TSA with 150 volume averages can visualize cell bodies near the ganglion cell layer (GCL) in the mouse retina [22]. Further, Pi *et al*. observed cell bodies appearing near the GCL of the rat retina, after 100 volume average, with a size distribution similar to RGCs [23]. These findings suggest that visualizing transparent RGC soma is possible using TSA. Future studies using TSA to track the distribution of RGCs in the tree shrew retina after the onset of optic neuropathy could provide insight into the rate of RGC loss with disease progression. Further, this rate could be correlated with the size reduction of individual axon bundles to better predict RGC soma density in the human retina.

RGC axon bundles consist of axonal membranes, microtubules, neurofilaments, and mitochondria [14, 29]. These bundles relay visual signals from RGC soma to the brain through the ONH [30]. Bundles passing through the ONH are organized such that bundles from the peripheral retina are on the outer sections of the optic nerve, while bundles nearest the ONH are on the inner portions [31]. Our fibergram images from the tree shrew retina revealed an RGC axon bundle pattern that was more similar to primates than rodents. In tree shrews, the bundles showed a higher density throughout most of the retina, which is consistent with the high bundle density in primates [32, 33]. By contrast, the rodent retina consists of bundles that form a sparsely distributed monolayer throughout the retina [5, 15, 16]. Similar to primates [33], tree shrew axon bundles in the periphery formed a monolayer that became more stratified as the bundles converged on the ONH. These stratifications consist of bundles that form in the periphery on the bottom-most of the RNFL, with bundles that form closer to the ONH located on the superficial part of the RNFL [31, 33]. Given that (1) the differentiation of RGCs begins in the central retina, more cells are added peripherally in a roughly concentric fashion [34-36], and (2) axons elongate on the inner surface of the retina, a favorable growth environment during early development [37, 38], axons from central retina will occupy the space close to the inner surface and axons from peripheral retina will be squeezed toward the deeper space in the RNFL. This is also observed in cross-sectional images of bundles in healthy rhesus monkey retinas [30, 32].

We quantified three parameters to define axon bundle sizes at different distances from the ONH in tree shrew retinas. For measurements taken throughout the whole retina, as bundles diverged from the ONH, width increased, and height decreased. However, the cross-sectional area of the bundles plateaued near 1.5 mm from the ONH. We analyzed a continuous 45° sector in one of the tree shrews. Throughout this sector, bundle width decreased towards the ONH, and height increased. The cross-sectional bundle area followed a less distinct trend within the sector, with a peak value at 1.5 mm from the ONH. As explained above, bundles are squeezed vertically close to the ONH. However, all bundles still need to touch the inner surface of the retina for axon elongation and pathway finding during early development [34-38]. Therefore, axon bundles from RGCs near the ONH form smaller bundles in the superficial portion of the RNFL, whereas the fully formed bundles from the periphery take the bottom-most part of the RNFL [32].

Given the greater degree of anatomical similarity between tree shrew eyes and human eyes, tree shrews are ideal for modeling the pathophysiology of optic neuropathies, including glaucoma. Samuels *et al*. induced OHT in the tree shrew using ferromagnetic beads to block the aqueous humor outflow pathway, resulting in RNFL thinning and axonal loss within the optic nerve [12]. In future studies, this tree shrew OHT model could be used to identify which RGC axon bundle size parameter is most sensitive to glaucoma progression.

